## Supplementary text and figures for "Conserved and Unique Features of Terminal Telomeric Sequences in ALT-Positive Cancer Cells"

**Legends to supplementary figures**

**Figure S1 Analysis of telomeric and centromeric reads and 5’ termini randomization. Related to Figure 1.**

**A.** Relative abundance of pan-centromeric and telomeric reads. RPM (reads per million) of pan-centromeric and telomeric reads found in the human T2T reference genome [48] . **B.** Normalized number of pan-centromeric and telomeric reads aligned to Watson and Crick strands in HeLa and RPE1-hTERT cell lines. For telomeres, 3 units of TTAGGG or CCCTAA were used to call Watson (green) or Crick (red) strands. For centromeres, 1 unit of AAACTAGACAGAAGCATT or AATGCTTCTGTCTAGTTT was used to call Watson (green) or Crick (red) strands. **C.** Sequence logo representing the conservation (bits) of the 5’ termini with absolute randomization (top) versus absolute precision (bottom). **D.** Effect of *in vitro* randomization of 5’ termini by T7 exonuclease. Replotting of data presented in Figure 1F. Percentage of telomeric reads that have the indicated sequence as a 5’ sequence that have as a 5’ end. **E-F.** Comparison of experimental data with modeled predictions to assess the impact of varying degrees of randomization on the CCAATC-5’ termini. The data presented in Figure 1F for in-plug treatment of T7 (E) and in Figure 1G for POT1 depletion (F) show varying degrees of randomization compared to their respective untreated controls. **G.** Relative expression level of POT1 in HeLa cells expression inducible shRNA against POT1 (shPOT1) compared to control (shScr) after 3 days of shRNA induction by doxycycline.

**Figure S2 CRISPR/Cas9-Mediated endogenous knock-in and knockout of POT1a and POT1b in mESCs with functional validation. Related to Figure 2.**

**A.** Schematic of the POT1a genomic locus (red) and repair template containing homology arms to the Pot1a locus (red), selection marker blasticidin (BSD), 3xflag tag and FKBP12^F36V^ (blue). Scissors represent the CRISPR/Cas9 cut site used for endogenous knock-in generation. Sanger sequencing conforms the newly generated FKBP12^F36V^  and *POT1a* junction. **B.** PCR genotyping to detect knock-in was performed. **C.** Western blot analysis of mESCs expressing Flagx3-FKBP12^F36V^::POT1a treated with or without dTAG13 (500 nM) for 24 hrs. **D.** Schematic of the POT1b genomic locus showing exons targeted by sgRNAs (exon 2, E2, in red and exon 4, E4, in blue) and untargeted exons (grey), with wild-type and knockout conditions indicated. Scissors mark the CRISPR/Cas9 cut site used to generate the knockout cell line. Primers for genotyping are labelled as P1, P2, and P3 with arrows. Sanger sequencing conforms the knock-out by generation of newly generated exon 2 and exon 4 junction, resulting in a deletion ad frameshift. **E.** PCR genotyping to detect knockdown was performed. **F.** Relative expression level of POT1b in mESCs compared to control. **G.** Comparison of experimental data presented in Fig 2A with modeled predictions to assess the impact of varying degrees of randomization on the CCAATC-5’ termini.

**Figure S3 Analysis of 5’ termini and of presence of variant telomeric repeats in U2OS. Related to Figure 3.**

**A-C.** Distribution of telomeric reads across biological replicates of ALT-positive G292 (A), SAOS2 (B) and U2OS cells. The following sequences (CCCAAT-5’, TCCCAA-5’, and ATCCCA-5’) are grouped and labelled as “Rest”. **D.** Relative expression level of POT1 in U2OS cells expression of two different inducible shRNA against POT1 (shPOT1-1 and shPOT1-2) compared to control (shScr) after 3 days of shRNA induction by doxycycline. **E.** Percentage of canonical telomeric reads (CTRs) and variant telomeric reads (VTRs) within the first 30 nucleotides from the 5’ of the terminal of Illumina reads from U2OS or HeLa cells.

**Figure S4 ssDNA in ALT telomeres. Related to Figure 4.**

**A-B.** Representative images and quantification of the percentage of ALT-positive U2OS cells and ALT-negative HeLa cells with more than 5 RPA foci colocalizing to TRF2. Data shown repeated in triplicate with a minimum of 250 cells counted per cell line. Error bars represent the standard error of the mean where n=3. The scale bar represents 10 μm. **C-D.** Representative images and quantification of the percentage of ALT-positive U2OS cells and ALT-negative HeLa cells with more than 5 FISH foci where the staining is done under native conditions (ssTelo) to label single-stranded G-rich telomeric DNA (ssTeloG, green) or C-rich telomeric DNA (ssTeloC, red). Data shown repeated in triplicate with a minimum of 250 cells counted per cell line. Error bars represent the standard error of the mean where n=3. The scale bar represents 10 μm. **E.** Number of reads containing at least 4 consecutive telomeric repeats corresponding to the G-rich (green) and C-rich (red) strands in a panel of ALT-negative and ALT-positive cells. Telomeric reads are normalized to the total number of reads identified by S1-END-seq and are represented as reads per million (RPM). **F.** Schematics representing the expected number of C-strand and G-strand reads obtained by S1-END-seq in the absence of ssDNA regions within telomeres (1 end), or in the presence of 1 or 2 ssDNA. Bottom graph represents the expected ratio of C/G reads normalized to the total number of telomeric reads. **G**. Table representing the expected number of C-strand and G-strand reads following a S1-END-seq in the presence of the listed number of ssDNA region with a single chromosome end. **H.** **.** Percentage of canonical telomeric reads (CTRs) and variant telomeric reads (VTRs) within the first 30 nucleotides from the 5’ of the terminal of Illumina reads from END-seq and S1-END-seq libraries of U2OS cells. **I.** Percentage of telomeric reads that have the indicated sequence as a 5’ sequence that have as a 5’ end. The following sequences (CCCAAT-5’, TCCCAA-5’, and ATCCCA-5’) are grouped and labelled as “Rest”. Cells expressing either a non-targeting shRNA (shScr) or a shRNA targeting POT1 (shPOT1-1 and shPOT1-2) were harvested 3 days post induction and analyzed by S1-END-seq. Kullback-Leibler divergence (KL divergence) analysis was used to compare the distributions of the individual conditions. KL divergence smaller than 0.125 are considered non-significant and indicated as n/s

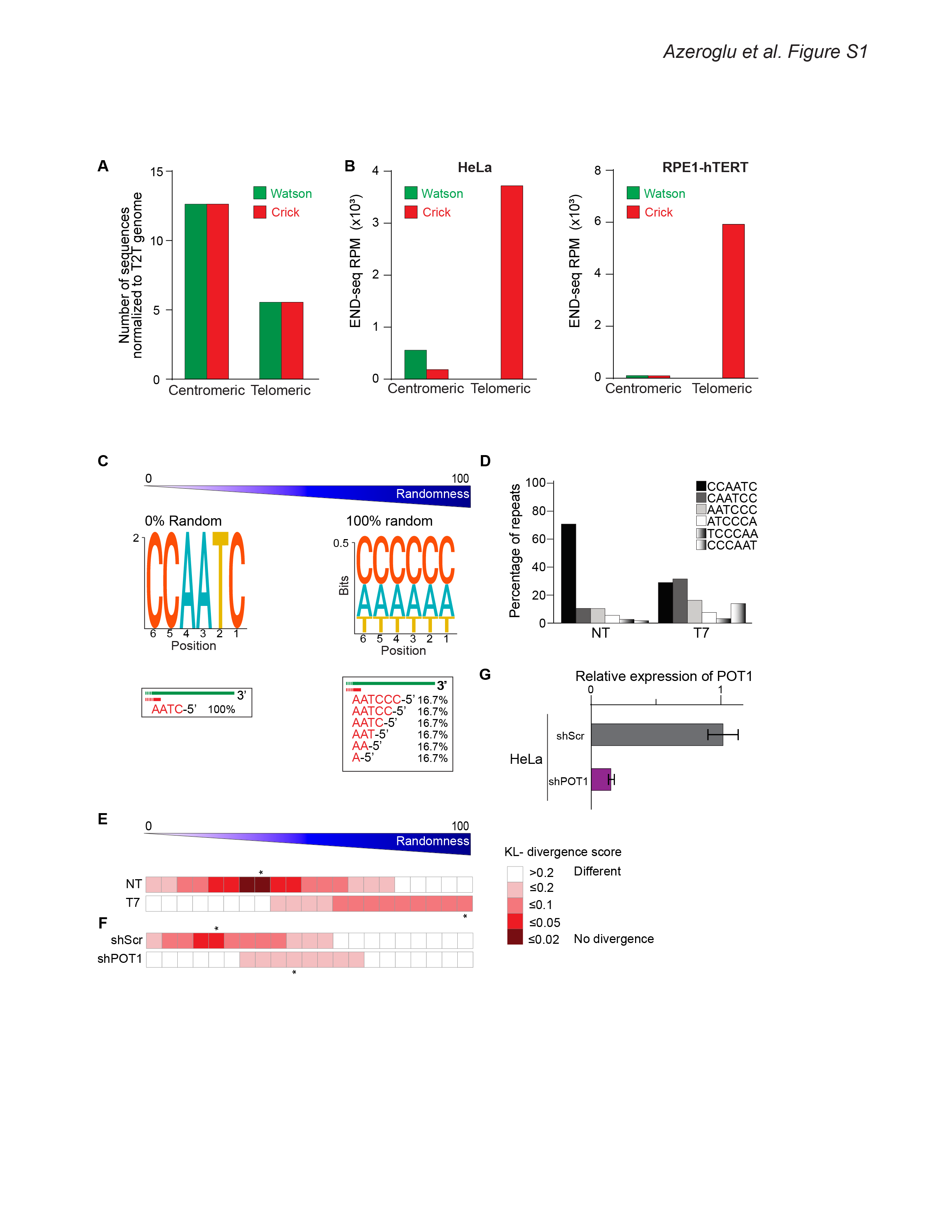

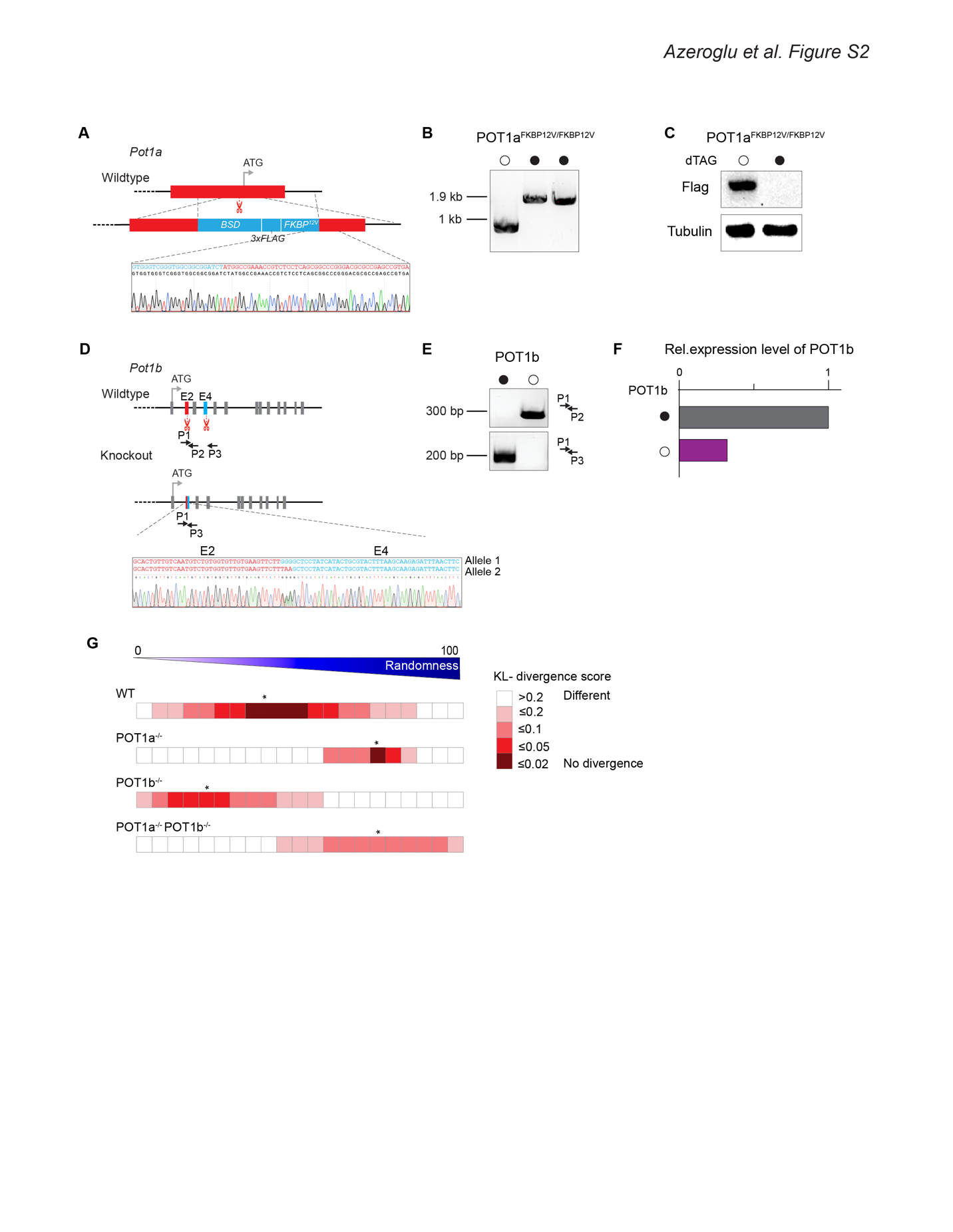

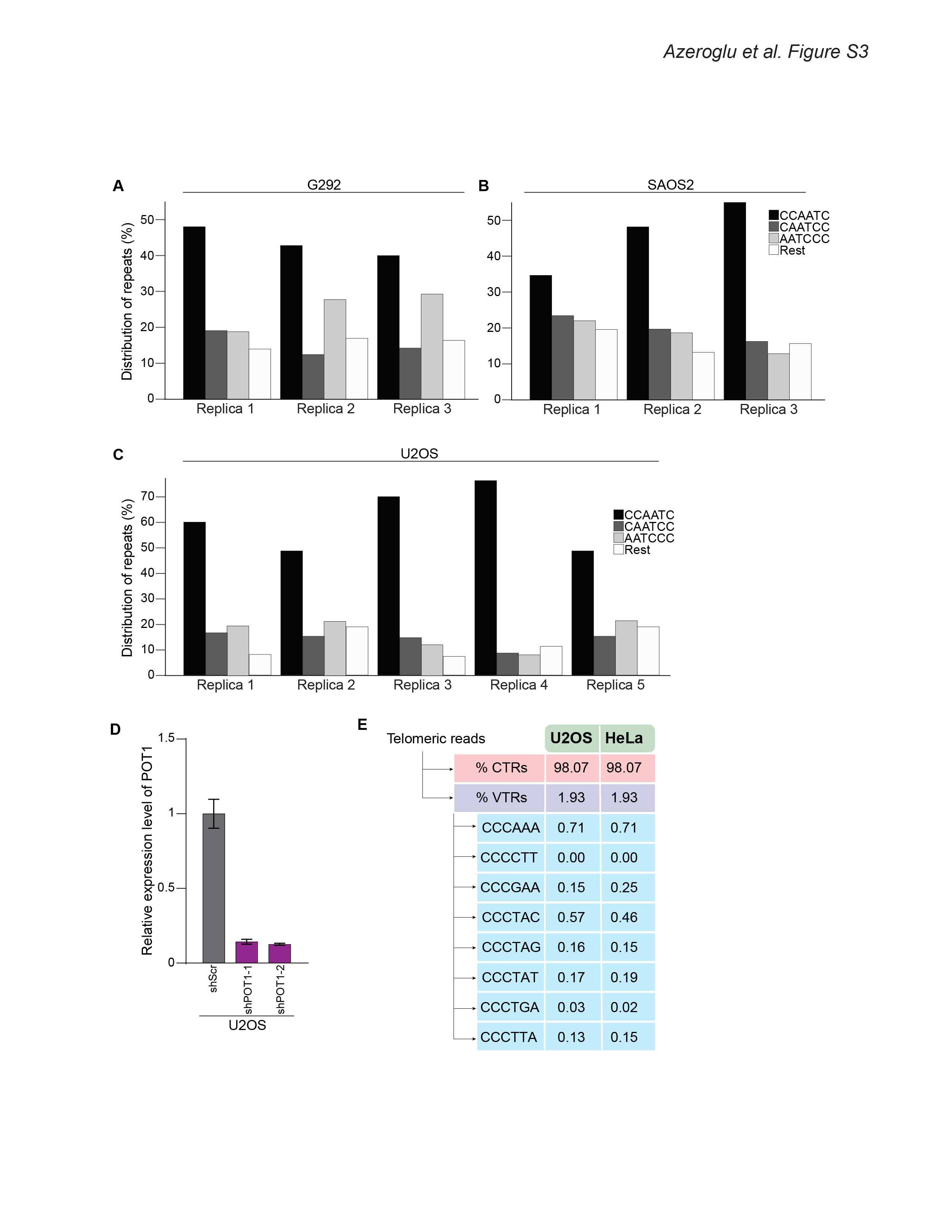

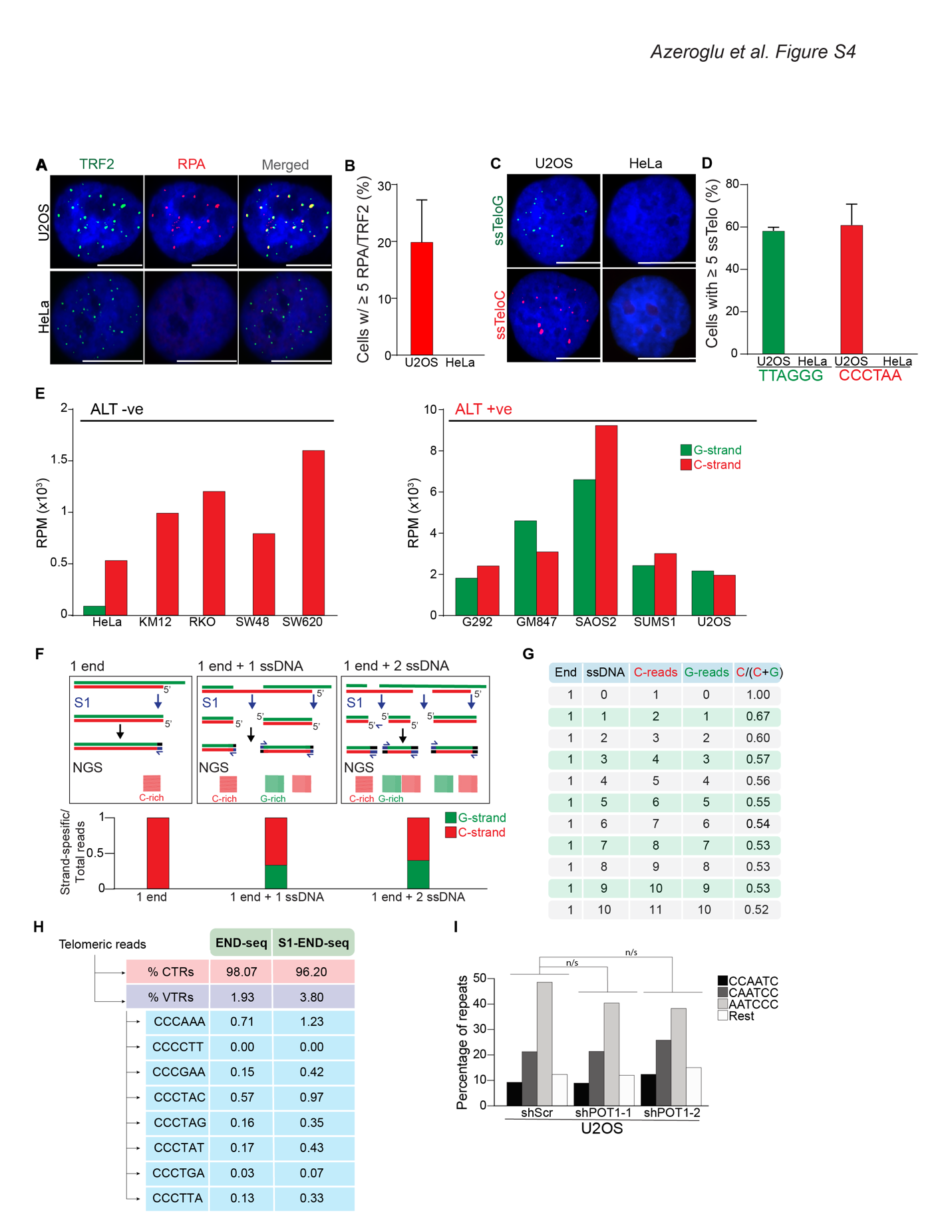
